## Supplementary material for "A Novel Mouse Model for *LAMA2*-Related Muscular Dystrophy: Analysis of Molecular Pathogenesis and Clinical Phenotype": Figure supplement, Table supplement: Supplementary Materials-3.12.pdf

### An Innovative Knockout Mouse Model for LAMA2-Congenital Muscular Dystrophy: Disruption of the Blood-Brain Barrier and Compromised Muscle Cytoskeleton Result in a Decreased Lifespan

Dandan Tan[1, 7], Yidan Liu[1], Huaxia Luo[1], Qiang Shen[2], Xingbo Long[3], Luzheng Xu[4], Jieyu Liu[1], Nanbert Zhong[5,*], Hong Zhang[2,*] & Hui Xiong[1, 6,*]

***Correspondence**

Hui Xiong, Department of Pediatrics, Peking University First Hospital, Beijing, 100034, China. Tel.: +8610-83573238;

Hong Zhang, Institute of Cardiovascular Sciences and Key Laboratory of Molecular Cardiovascular Sciences, Peking University Health Science Center, Beijing, 100191, China. Tel.: +8610-82801510;

Nanbert Zhong, New York State Institute for Basic Research in Developmental Disabilities, 1050 Forest Hill Road, Staten Island, NY 10314, USA. Tel/Fax: +1 718 494 5242/4882;

#### Figure supplement legends

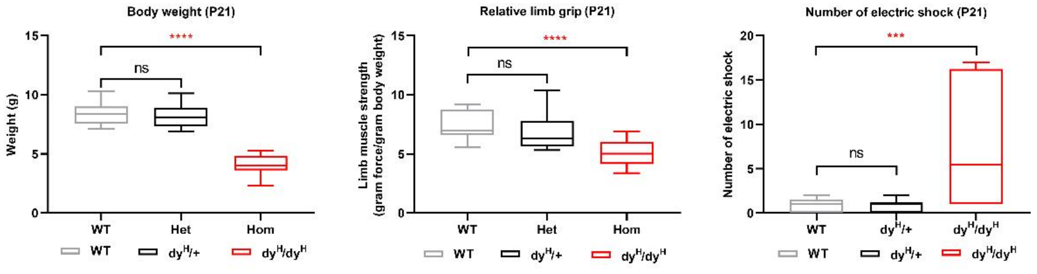


**Figure supplement 1. Body weight and muscle function analysis of $dy^H/dy^H$, WT and Het mice.** Significant differences were showed in body weight, the mean relative four-limb grip (force per gram body weight), and the number of electric shocks on the treadmill between the WT and $dy^H/dy^H$ mice at P21.

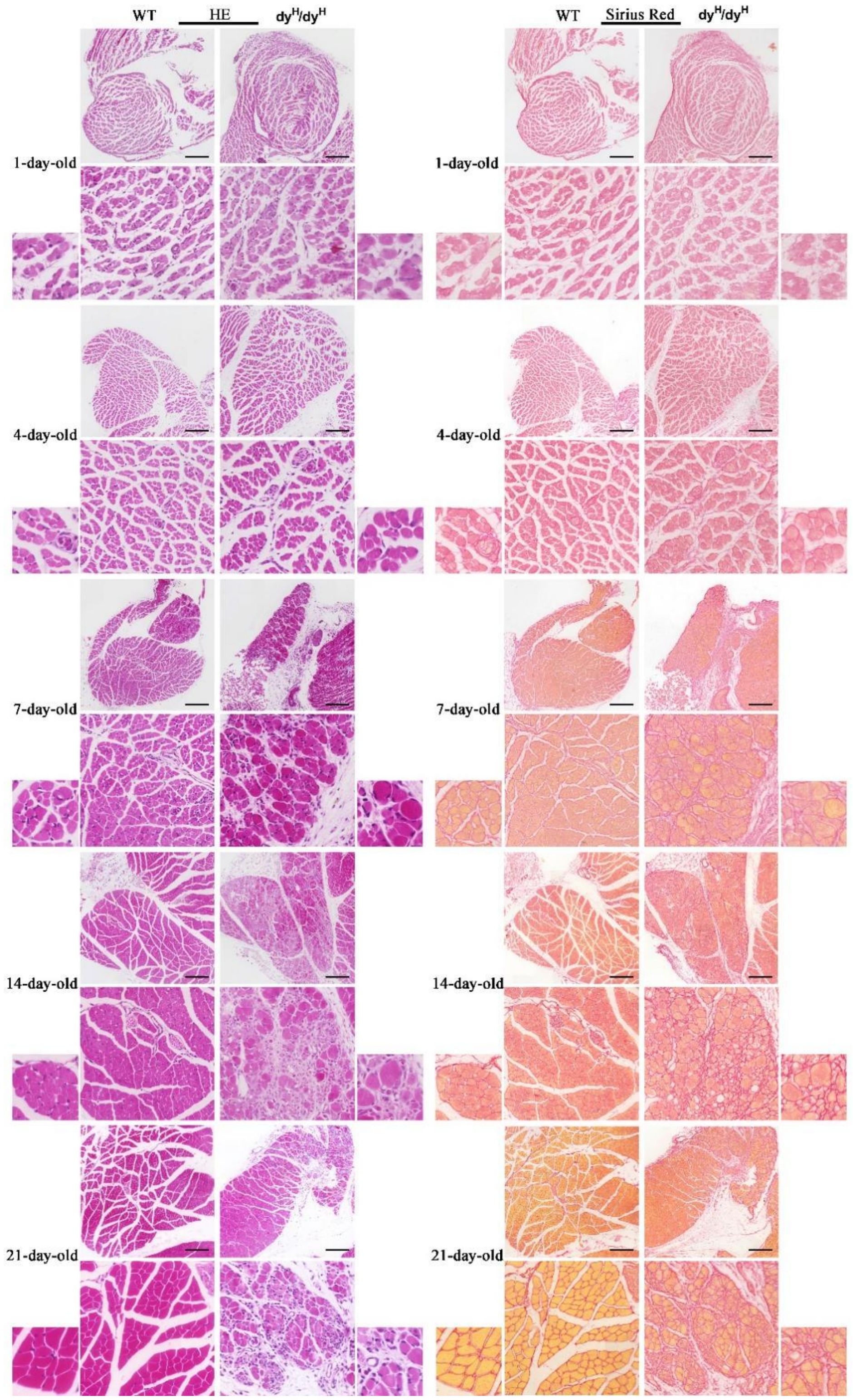


**Figure supplement 2. Muscle pathology with age in the biceps femoris of $dy^H/dy^H$ mice.** H&E and Sirius Red staining of the biceps femoris were compared between wild-type and $dy^H/dy^H$ mice at P1, P4, P7, P14 and P21. Whole biceps femoris, representative muscle area and magnifed panels were shown. Scale bars: for the whole muscle: 200 μm; intermediate magnifcation panel: 50 μm, maximum magnifcation panel: 20 μm.

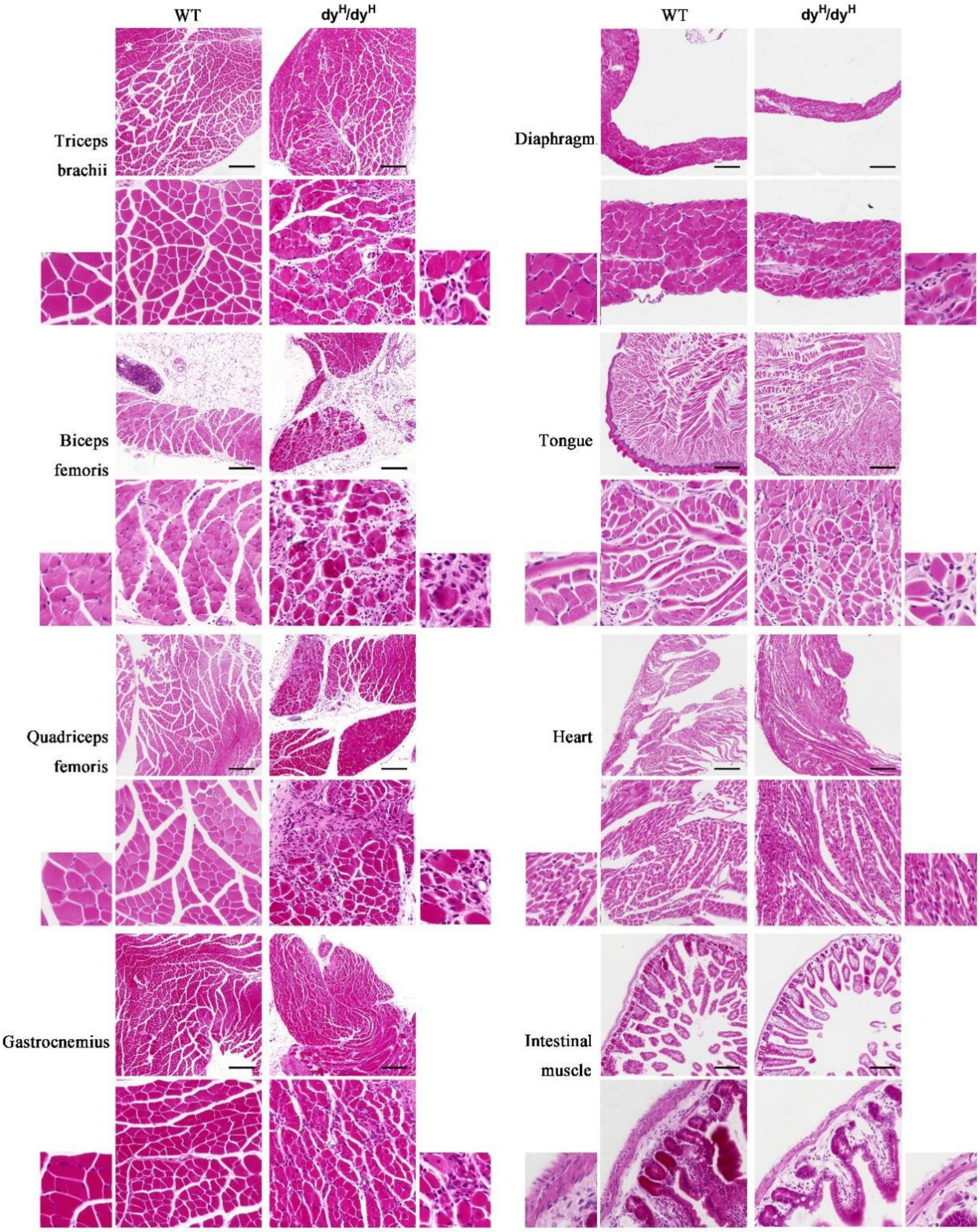


**Figure supplement 3. Extensive involvement with muscle pathology in dy$^{H}$/dy$^{H}$ mice at P21.** H&E staining showed dystrophic changes in the quadriceps femoris, gastrocnemius, triceps brachii, diaphragm and tongue muscles, but not as much as in the biceps femoris, while the heart and intestinal muscles were spared. Whole muscle, representative muscle area and magnifed panels were shown. Scale bars: for the whole muscle: 200 µm; intermediate magnifcation panel: 50 µm, maximum magnifcation panel: 20 µm.

A

| Sample | Total number of cells | Total number of cells after filtering | Mean Reads per Cell | Median Genes per Cell | Total Genes Detected |
|---|---|---|---|---|---|
| $dy^{H}/dy^{H}$ brain | 11,039 | 10,094 | 37,171 | 2,222 | 23,124 |
| WT brain | 10,540 | 10,496 | 38,995 | 2,080 | 22,964 |

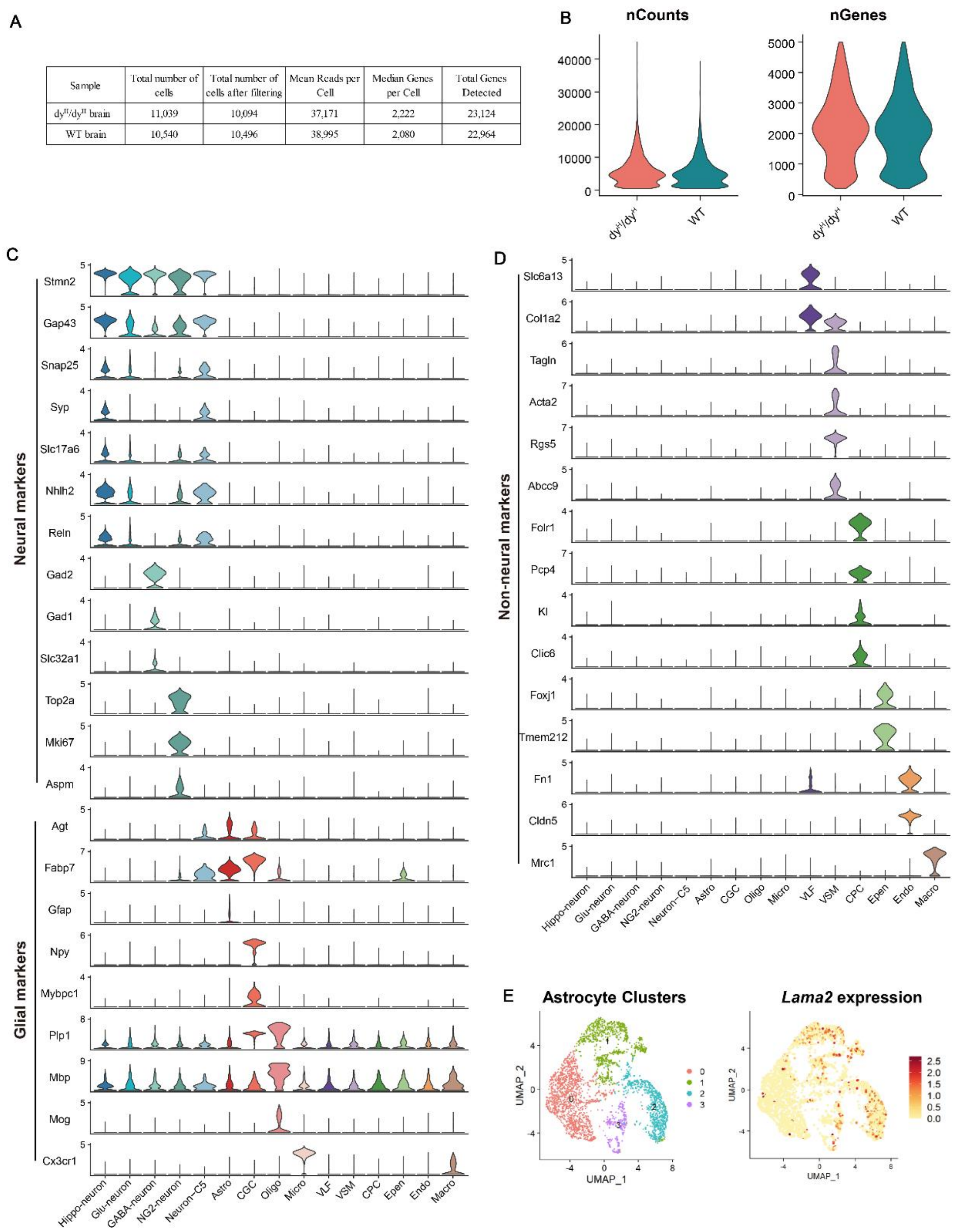


**Figure supplement 4. scRNA-seq analysis of $dy^{H}/dy^{H}$ and WT mouse brains. (A)** Statistics of the sequencing results from each sample. **(B)** Violin plots show the number of UMI counts and detected genes in each sample. **(C, D)** Violin plots of the expression distribution of the selected markers for each cell clusters. The cell cluster annotation and the corresponding markers are as follow. neuron: Stmn2, Gap43, Snap25, Syp; hippocampal neuron (Hippo-neuron): Nhlh2, Reln; glutamatergic neuron (Glu-neuron): Slc17a6; GABAergic neuron (GABA-neuron): Gad2, Gad1, Slc32a1; neuron-glial antigen 2 neuron (NG2-neuron): Top2a, Mki67, Aspm; sstrocyte (Aso): Agt, Fabp7, Gfap; cerebellum glia cell (CGC): Npy, Mybpc1; oligodendrocyte (Oligo): Plp1, Mbp, Mog; microglia (Micro): Cx3cr1; vascular and leptomeningeal fibroblasts (VLF): Slc6a13, Col1a2; vascular smooth muscle cell (VSM): Tagln, Acta2, Rgs5, Abcc9; choroid plexus cell (CPC): Folr1, Pcp4, Kl, Clic6; ependymal cell (Epen): Foxj1, Tmem212; endothelial cells (Endo): Fn1, Cldn5; macrophage (Macro): Mrc1. **(E)** UMAP visualization of *Lama2* expression in astrocyte clusters.

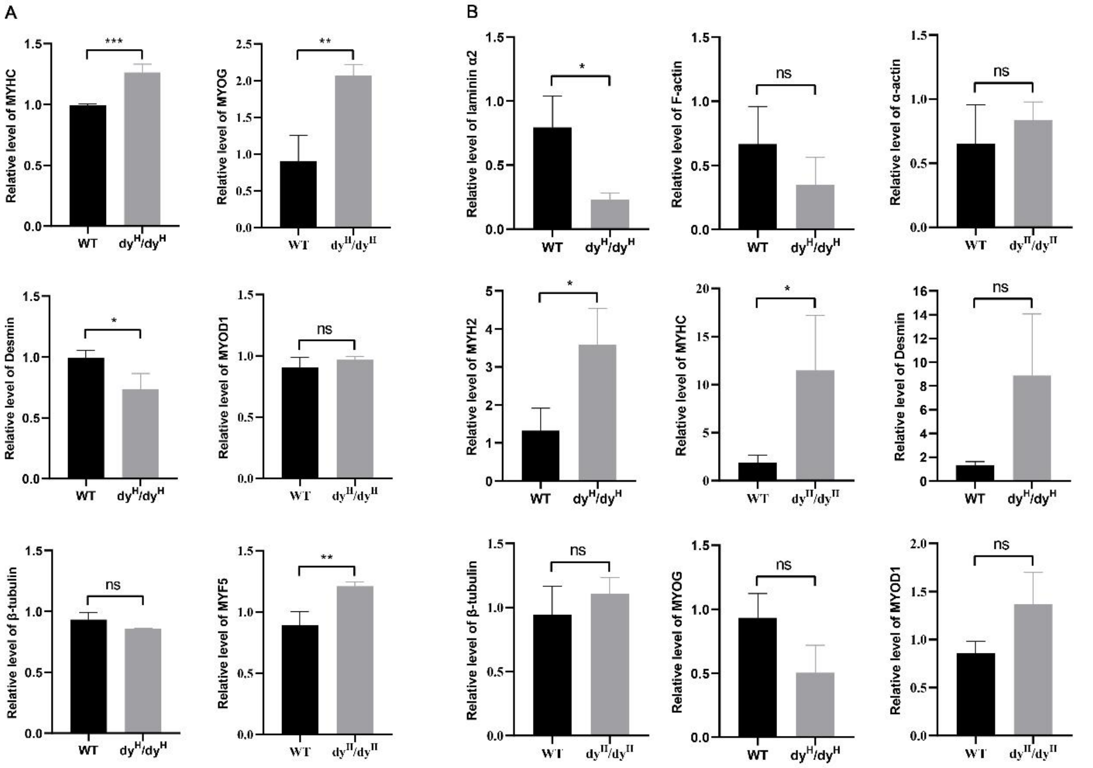


**Figure supplement 5. Quantitative analysis of muscle cytoskeleton and development proteins in $dy^H/dy^H$ mice. (A)** Quantitative analysis of immunofluorescence staining for MYHC, MYOG, desmin, MYOD1, β-tubulin, and MYF-5 in WT and $dy^H/dy^H$ muscles at P14. **(B)** Quantitative analysis of Western blot for laminin α2, F-actin, α-actin, MYH2, MYHC, desmin, β-tubulin, MYOG and MYOD1 in P14 WT and $dy^H/dy^H$ muscles. *** for p-value< 0.001, ** for p-value<0.01, * for p-value<0.05.

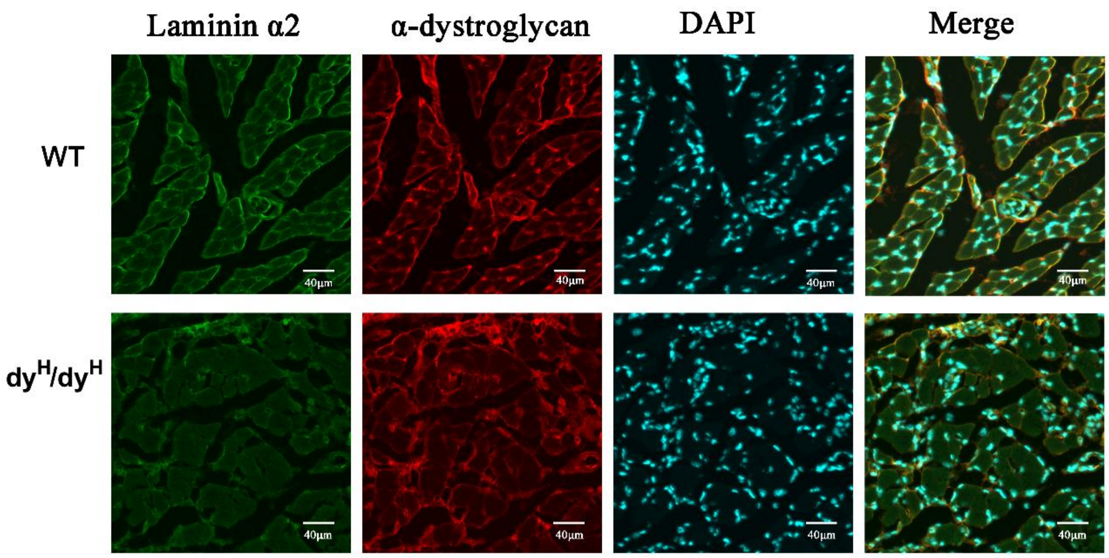


**Figure supplement 6. Immunofluorescence staining for α-dystroglycan in the biceps femoris of $dy^H/dy^H$ mice.** Colocalization of α-dystroglycan (red fluorescence, blue arrow) and laminin α2 chain (green fluorescence, yellow arrow) showed that α-dystroglycan protein was normally located and expressed in the cell membrane both in the WT and $dy^H/dy^H$ muscles. Scale bars: 40 μm.

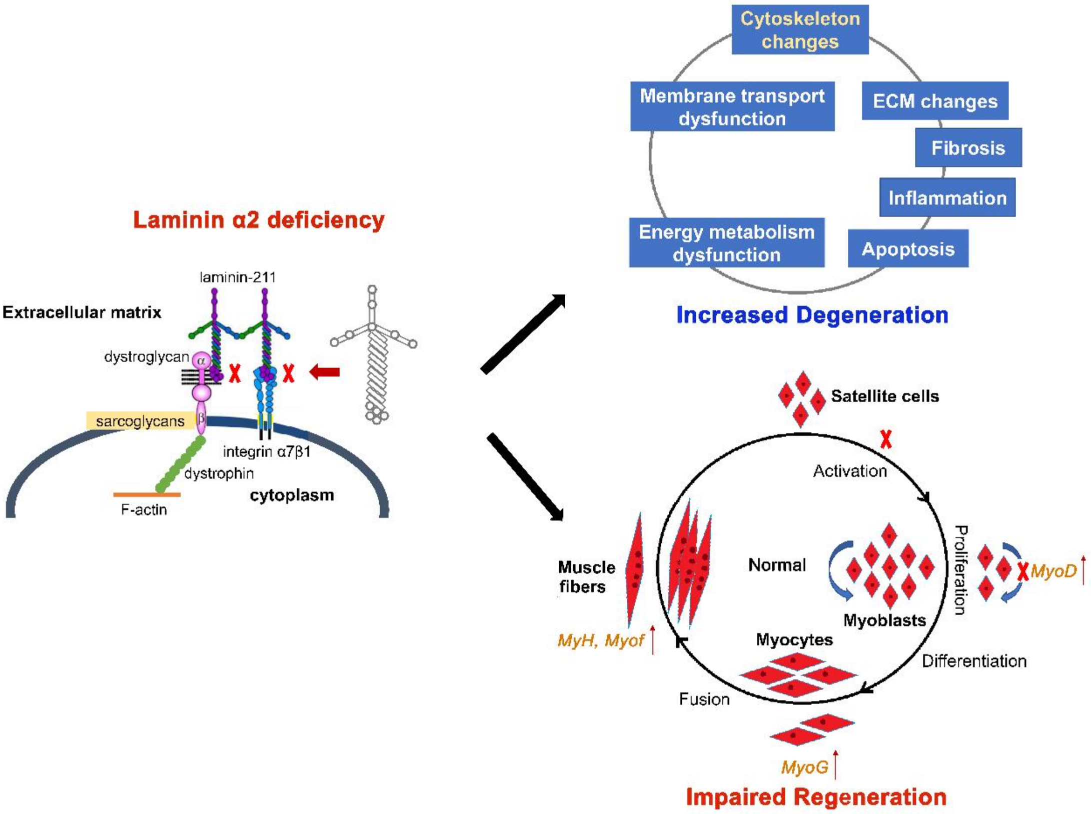


**Figure supplement 7. Proposed pathogenic mechanisms hypothesis in the dy$^{H}$/dy$^{H}$ mouse model of *LAMA2*-CMD.** Laminin α2 deficiency in the basement membrane due to pathogenic variants in the *LAMA2* gene leads to loss of dystroglycan/integrin-matrix scaffolds and extracellular matrix networks. Then, a series of molecular changes associated with the cytoskeleton, extracellular matrix, fibrosis, inflammation, apoptosis, pyroptosis, and mitochondrial energy metabolism occur.

**Table supplement 1. Differential expressed genes in all cell clusters in scRNA-seq analysis.**

**Table supplement 2. Differentially expressed genes (DEGs) (1136 upregulated and 884 downregulated) at least two-fold ($p$ < 0.05) in dy$^{H}$/dy$^{H}$ (KO) muscle samples relative to the wildtype (WT) muscle samples.**

**Table supplement 3. Comparison between the $dy^H/dy^H$ mouse with other *Lama2* deficient mice.**

| Mouse | *Lama2* mutaion | Laminin α2 expression | Muscular dystrophy | BBB Deficits | Life expectancy | References |
|---|---|---|---|---|---|---|
| $dy^H/dy^H$ | Knock-out by out-frame deletion of the exon 3 | Complete deficiency | Very severe | BBB dysfunction with laminin α2 deficiency | 3 weeks of age | This paper |
| dy/dy | Spontaneous, unknown | Reduced expression of normal sized laminin α2 | Moderate | Unknown | Before 6 months of age | *Xu et al., 1994; Michelson et al., 1995* |
| $dy^{2J}/dy^{2J}$ | Spontaneous splice site mutation resulting in an in-frame deletion in the exon 2 | Slightly reduced expression of truncated laminin α2 | Mild | Unknown | After 6 months of age | *Sunada et al., 1995* |
| $dy^{3k}/dy^{3k}$ | Knock-out by inserting a reverse Neo element in the 3' end of exon 4 | Complete deficiency | Very severe | BBB dysfunction and increased permeability | 3 weeks of age | *Miyagoe et al., 1997;* Gawlik *et al., 2019; Menezes et al., 2014* |
| $dy^W/dy^W$ | Knock-out by inserting lacZ-neo element in the exon 1 | Severely reduced expression of truncated laminin α2 | Severe | Unknown | 5–12 weeks of age | *Kuang et al., 1998* |

*Abbreviations:* BBB, blood-brain barrier.

**Table supplement 4. Primers of PCR-amplifications for Genotype identification.**

| Primers | Primers' sequence (5'-3') | Lengths of PCR products (bp) |
|---|---|---|
| *Lama2*-F | ACTGAACCCAGGCTCCCTTTGAATC | wild-type: 571 |
| *Lama2*-R1 | ATTAGACATCGAACCACCTCTGTTTTCA | mutant-type: 0 |
| *Lama2*-F | ACTGAACCCAGGCTCCCTTTGAATC | wild-type: 1999 |
| *Lama2*-R2 | AACCTCAAGGCTGACACCCTGCTAA | mutant-type: 374 |

*Abbreviations:* PCR, polymerase chain reaction.

**Table supplement 5. MRI examination imaging sequences.**

| Series description | Plane | TR (ms) | TE (ms) | FOV (mm) | Slice thickness (mm) | Distance factor (%) | Matrix | BW (Hz/pixel) |
|---|---|---|---|---|---|---|---|---|
| T2-weithted (TSE) | Coronal | 3000 | 78 | 35 | 1 | 10 | 256×256 | 174 |
| T1-weighted (SE) | Coronal | 250 | 16 | 35 | 1 | 10 | 256×256 | 120 |

*Abbreviations:* TR, time of repetition; TE, time of echo; FOV, field of view; BW, bandwidth; TSE, turbo spin echo; SE, spin echo.
